# Skill learning can be independent of speed and accuracy instructions

**DOI:** 10.1101/726315

**Authors:** Teodóra Vékony, Hanna Marossy, Anita Must, László Vécsei, Karolina Janacsek, Dezso Nemeth

## Abstract

A crucial question in skill learning research is how instruction affects the performance or the underlying representations. However, a little is known about its effect on one critical aspect of skill leaning, namely, picking-up statistical regularities. More specifically, how pre-learning speed vs. accuracy instructions affect the acquisition of non-adjacent second-order dependencies. Here, we trained two groups of participants on an implicit probabilistic sequence learning task: one group focusing on being fast and the other on being accurate. As expected, we detected strong instruction effect: accuracy instruction resulted in a nearly errorless performance, while speed instruction caused short reaction times. Despite the differences in the average reaction times and accuracy scores, we found a similar level of statistical learning in the training phase. After the training phase, we tested the two groups under the same instruction (focusing on both speed and accuracy), and they showed comparable performance, suggesting a similar level of underlying statistical representations. Our findings support that skill learning can result in robust representations, and they highlight that this form of knowledge may appear with almost errorless performance.

## Introduction

Our social, motor and cognitive skills help us to adapt and function in various situations in our everyday life. Therefore, our ability to learn new skills is crucial, and fine-tuning this capability can be advantageous for an individual. Previous studies investigating sports performance (Beilock, Bertenthal, Hoerger, & Carr, 2008; Beilock, Bertenthal, McCoy, & Carr, 2004), implicit (Hoyndorf & Haider, 2009), and explicit sequence learning (Barnhoorn, Panzer, Godde, & Verwey, 2019) typically found that speed and accuracy strategies differently affects skill learning. However, skill learning is multifaceted, and it still not clear what underlying mechanisms benefit from speed and accuracy instructions and what mechanisms do not. A core component of learning new skills is picking up complex statistical regularities from the environment (Conway, 2020; Janacsek, Fiser, & Nemeth, 2012). So far, no study has investigated the effects of prioritizing speed or accuracy on the acquisition of such statistical dependencies. Do such instructions affect our performance or also the established statistical representations? Above the theoretical importance of answering this question, it also has clear practical implications. If instructions do affect statistical knowledge acquisition itself, then differences in the exact instruction between studies or the interpretation of instructions among participants might affect the conclusions about skill learning. Here, we aim to unveil how emphasizing speed and accuracy influence an essential aspect of skill learning, namely the acquisition of complex statistical representations.

Hoyndorf and Haider (2009) investigated the sequencing aspect of skill learning and found accuracy strategy to impair the expression of implicit knowledge compared to speed instruction; however, learning was still detected under accuracy instruction compared to a non-learning control group. Yet in this experiment, the accumulated sequence-knowledge under speed/accuracy instructions was not compared to a phase where the importance of speed and accuracy was equally emphasized. Such a comparison would reveal whether implicit sequence knowledge is acquired to the same level under different instructions. Recently, Barnhoorn, Panzer, Godde, and Verwey (2019) found that speed instruction benefits the development of representations about repeating sequences while forcing participants to be more accurate leads to faster selection of responses via better stimulus-response associations. In this study, the participants were aware of the repeating sequences; thus the learning was utterly explicit. Taken together, the studies mentioned above suggest that speed instruction might benefit sequence learning more than accuracy instruction. These studies used relatively simple, deterministic sequences (i.e., sequences with a simple repeating pattern). Therefore, data are still lacking on whether instruction affects more complex, probabilistic sequence representations.

Here, we aimed to test whether speed or accuracy instructions affect the acquisition of complex statistical regularities using an implicit probabilistic sequence learning task. We go beyond previous investigations by at least two aspects: First, by studying complex probabilistic sequences with non-adjacent, second-order dependencies (Remillard, 2008). This feature means that to predict the n^th^ element of the sequence, we need to know the n-2^th^ element instead of n-1^th^. This structure creates an abstract sequence representation, and its acquisition will be based on statistical regularities (Nemeth et al., 2013), which are also fundamental in complex cognitive skills such as the human language (Christiansen & Chater, 2015). The second novel contribution of our study is that we also test the implicit sequence knowledge of our participants a*fter* the (instructed) training phase. Our learning task was completed in two different phases. In the first phase, we instructed the participants to focus either on accuracy or speed while performing the task (Different Instruction Phase, Accuracy vs. Speed Group). After the training phase, we tested both groups of participants with the same instruction (i.e., focusing *both* on accuracy and speed, Similar Instruction Phase). By doing so, we aimed to differentiate between the effects of instructions on the training performance and the acquired knowledge. Our questions were 1) whether the speed/accuracy instruction affects the learning of probabilistic statistical regularities; and if instructions do modify the learning process 2) do they affect the training performance (Different Instruction Phase) and the retrieval of knowledge (Similar Instruction Phase) equally?

## Materials and Methods

### Participants

Sixty-six healthy young adults took part in the study. Five of them were excluded from the experiment because they conceivably misunderstood the instructions. Their performance was more than two standard deviations from the mean of their group in more than 50% of the epochs (units of analysis), which was not observable during the practice session. Therefore, 61 participants remained in the final sample (40 females). They were between 19 and 27 years of age (*M*_*age*_ = 21.18 years, *SD*_*age*_ = 2.13 years). All of them were undergraduate students from Budapest, Hungary (*M*_*years of education*_ = 14.14 years, *SD*_*years of education*_ = 1.64 years). Participants had normal or corrected-to-normal vision, none of them reported a history of any neurological and/or psychiatric disorders, and none of them was taking any psychoactive medication at the time of the experiment. Handedness was measured by the Edinburgh Handedness Inventory (Oldfield, 1971). The Laterality Quotient (LQ) of the sample varied between -84.62 and 100 (−100 means complete left-handedness, 100 means complete right-handedness; *M*_*LQ*_ = 62.25, *SD*_*LQ*_ = 53.73). They performed in the normal range on the Counting Span Task (*M*_*Counting Span*_ = 3.66, *SD*_*Counting Span*_ = 0.81) All of the participants gave written informed consent before enrollment and received course credit for participating. They were randomly assigned to the Accuracy Group (n = 31) or Speed Group (n = 30). No group differences were observed in terms of age, years of education, handedness, and neuropsychological performance (see Table 1). The study was approved by the Research Ethics Committee of the Eötvös Loránd University, Budapest, Hungary, and it was conducted in accordance with the Declaration of Helsinki.

**Table 1.** Comparison of the two groups on age, years of education, handedness and neuropsychological performance

|  | *Accuracy Group* | *Speed Group* | *t-test* |
| --- | --- | --- | --- |
|  | *M(SD)* | *M(SD)* |  |
| Age (years) | 21.29 (2.28) | 21.07 (2.00) | $t(59) = 0.41, p = .69$ |
| Education (years) | 14.31 (1.60) | 13.97 (1.71) | $t(59) = 0.80, p = .43$ |
| Handedness (LQ) | 54.88 (55) | 69.86 (52.2) | $t(59) = -1.09, p = .28$ |
| Counting Span Score | 3.69 (0.75) | 3.64 (0.88) | $t(59) = 0.21, p = .83$ |

### Alternating Serial Reaction Time task

In this study, we used the implicit version of the Alternating Serial Reaction Time (ASRT) task (J. H. Howard & Howard, 1997; Nemeth, Janacsek, Londe, et al., 2010). In the ASRT task, four empty circles (300 pixels each) were presented horizontally in front of a white background in the middle of a computer screen. A target stimulus (a drawing of a dog’s head, 300 pixels) was presented sequentially in one of the four empty circles (Figure 1A). A keyboard with four heightened keys (Z, C, B, and M on a QWERTY keyboard) was used as a response device, each of the four keys corresponding to the circles in a horizontal arrangement. Participants were asked to respond with their middle and index fingers of both hands by pressing the button corresponding to the target position. At the beginning of each block of the ASRT task, the four empty circles appeared horizontally on the screen for 200 ms, and then, the first target stimulus occurred, and it remained on the screen until the first correct response. The next stimulus appeared after a 120 ms response-to-stimulus interval.

**Figure 1.**
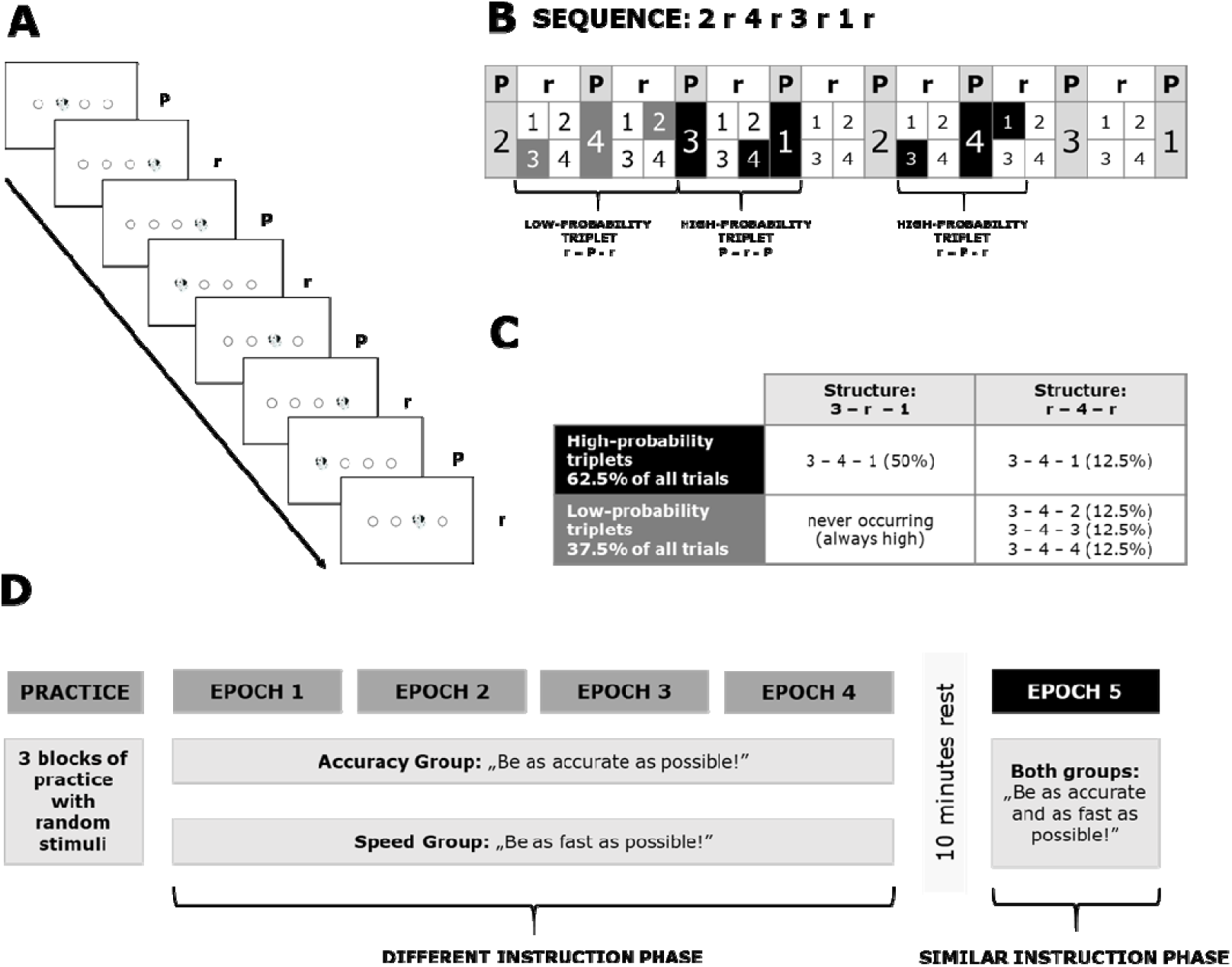
Task and design of the experiment. (A) Stimulus presentation in the ASRT task. A dog’s head appeared in one of the four positions. Stimuli appeared either in a pattern (P) or a random (r) position, creating an eight-item long alternating sequence structure. (B) High- and low-probability triplets. Due to the alternating sequence structure, some runs of consecutive visual stimuli (called *triplets*) occurred with a higher probability than others. Every trial was defined as the third trial of a high- or a low-probability triplet, based on the two preceding trials. High-probability triplets can be formed by two patterns and one random element, but also by two random and one pattern element. (C) The proportion of high- and low-probability triplets. High-probability triplets occurred in 62.5% of all trials (of which 50% came from pattern trials, i.e., from P-r-P structure, and 12.5% came from random trials, i.e., from r-P-r structure, by chance). Low-probability triplets occurred in the remaining 37.5% of all trials (of which each individual low-probability triplet occurred with a 12.5% probability by chance, originating only from r-P-r structure). (D) The design of the study. In the Different Instruction Phase, different instruction was told to the Accuracy and the Speed Group. After four epochs (each containing five blocks) of the ASRT task, and a 10-minute long rest period, the instruction changed. In the fifth epoch (containing five blocks of stimuli), the same instruction was given to all of the participants (Similar Instruction Phase).

The serial order of the four possible positions (coded as 1, 2, 3, and 4) in which target stimuli could appear was determined by an eight-element probabilistic sequence. In this 6 sequence, every second element appeared in the same order, while the other elements’ positions were randomly chosen out of the four possible locations (e.g., 2r4r3r1r; where r indicates a random position). Therefore, some combinations of three consecutive trials (*triplets*) occur with a greater probability than others. For example, 2_**4**, 4_**3**, 3_**1**, and 1_**2** (where ‘‘_” indicates any possible middle element of the triplet) would often occur because the third element (bold numbers) could be derived from the sequence (or occasionally could be a random element as well). In contrast, 1_**3** or 4_**2** would occur with less probability because the third element could only be random (Figure 1B). Therefore, the third element of a high-probability triplet is more predictable from the first event when compared to a low-probability triplet. There were 64 possible triplets in the task (four stimuli combined for three consecutive trials). Sixteen of them were high-probability triplets, each of them occurring in approximately 4% of the trials, about five times more often than the low-probability triplets. Overall, high-probability triplets occur with approximately 62.5% probability during the task, while low-probability triplets only occur with a probability of 37.5% (Figure 1C). As participants practice the ASRT task, their responses become faster and more accurate to the high-probability triplets compared to the low-probability triplets, revealing statistical learning throughout the task (J. H. Howard & Howard, 1997; Kóbor et al., 2017; Song, Howard, & Howard, 2007; Unoka et al., 2017). Each block of the ASRT task contained 85 stimuli (5 random elements at the beginning of the block, then the 8-element alternating sequence repeated 10 times).

### Inclusion-Exclusion Task

We also administered the Inclusion-Exclusion Task (Destrebecqz & Cleeremans, 2001; Destrebecqz et al., 2005; Fu, Dienes, & Fu, 2010; Jiménez, Vaquero, & Lupiáñez, 2006), which is based on the “Process Dissociation Procedure” (Jacoby, 1991). In the first part of the task, we asked participants in what order the stimuli (both pattern and random elements) appeared during the task, and they were asked to type the sequence using the same four response buttons the participants used during the ASRT task (inclusion instruction). After that, they had to generate new sequences that are different from the learned one (exclusion condition). Both parts consisted of four runs, and each run finished after 24 button presses, which is equal to three rounds of the eight-element alternating sequence (Horvath, Torok, Pesthy, Nemeth, & Janacsek, 2018; Kiss, Nemeth, & Janacsek, 2019; Kóbor et al., 2017). The successful performance in the inclusion condition can be achieved by solely implicit knowledge (however, explicit knowledge can also boost performance, but it is not necessary to the successful completion of the task). On the contrary, successful performance in the exclusion condition (i.e., generating a *new* sequence that is different from the learned one) can only occur if the participant has conscious knowledge about the learned statistical regularities. Generation of the learned statistical regularities above chance level even in the exclusion task indicates that the participant relies on their implicit knowledge, as it cannot be controlled consciously. To test whether the participants gained consciously accessible triplet knowledge, first, we calculated the percentage of the generated high-probability triplets in the inclusion and exclusion condition separately. Then we tested whether the occurrence of high-probability triplets differs from the probability of generating them by chance. We also compared the percentages of the high-probability triplets across conditions (inclusion and exclusion task) and groups (Accuracy Group and Speed Group) (for more details about the Inclusion-Exclusion task, see: Horvath et al., 2018; Kiss et al., 2019; Kobor et al., 2017).

### Questionnaire

We used a questionnaire to scrutinize whether the participants prefer accuracy or speed in general and whether they are rather accurate or fast in their everyday life. The questionnaire consisted of the following questions: “*In an everyday situation, what do you attend more: speed or accuracy (in a scale from 1 to 10, where 1 means that only the accuracy is important and 10 means that only the speed is important)*?”, “*In an everyday situation, how important is for you to be accurate/fast in a scale from 1 to 10?*”, *“According to your friends and family, how fast/accurate are you when you need to solve a problem (in a scale from 1 to 10)?”*.

### Design

First, the participants completed three practice blocks of 85 random trials each to familiarize themselves with the task. After that, the participants completed two sessions of the ASRT task. In the first, training session (referred to as Different Instruction Phase), we gave different instructions to the participants of the Accuracy and Speed Group. For the Accuracy Group, the instruction was to try to be as accurate as possible during the task. On the contrary, the instruction for the Speed Group was to be as quick as possible. By this, we could investigate the performance on the task in light of the different instructions. Twenty blocks were presented to the participants in the Different Instruction Phase (for analysis, we organized the blocks into four epochs by merging five consecutive blocks). Participants could rest a bit after each block. A 10 min rest period was inserted before the second ASRT session. During this period, participants were not involved in any demanding cognitive activity. The second session of ASRT (referred to as Similar Instruction Phase) contained five blocks (one epoch). This time, both the Accuracy and Speed Group were instructed to respond to the target stimulus as quickly and as accurately as possible (Figure 1D). After the ASRT task, the Inclusion-Exclusion task was administered.

### Statistical analysis

We defined each trial as the third element of a high or low-probability triplet. Trills (e.g., 1-2-1) and repetitions (e.g., 1-1-1) were eliminated from the analysis because participants tend to show pre-existing response tendencies to these type of triplets (D. V. Howard et al., 2004; Janacsek, Borbély-Ipkovich, Nemeth, & Gonda, 2018; Takács et al., 2018; Unoka et al., 2017). The first five button presses were random; thus, only the eighth button press could be evaluated as the last element of a valid triplet. Therefore, the first seven trials were excluded from the analysis. Blocks were collapsed into four epochs in the Different Instruction Phase (Epoch 1-4), and one epoch in the Similar Instruction Phase (Epoch 5) to facilitate data processing and to reduce intra-individual variability. We calculated the median reaction times (RTs) separately for high- and low-probability triplets for each participant and each epoch. Only correct responses were considered for the RT analysis.

We used mixed-design ANOVAs to compare the learning performance between the two groups in the Different and Similar Instruction Phase. In the Different Instruction Phase, we included the factors of Epoch (Epoch 1 to 4), Triplet (high- vs. low-probability triplets), and Group (Accuracy Group vs. Speed Group). The same analyses were performed for the Similar Instruction Phase, except that here the Epoch factor was not considered (as only one epoch was performed in this phase). In all ANOVAs, the Greenhouse-Geisser epsilon (ε) correction was used if necessary. Corrected *df* values and corrected *p* values are reported (if applicable) along with partial eta-squared (η_p_^2^) as the measure of effect size. We used LSD (Least Significant Difference) tests for pair-wise comparisons. We calculated Bayes-factors (BF) for the support of our non-significant, but relevant group comparisons. All of the frequentist analysis was carried out by using IBM SPSS Statistics 25, and all of the Bayesian analyses in JASP (JASP Team, 2019).

The instructions about the accuracy and speed during the experiment could cause major differences in the average RTs (i.e., all RTs to valid trials collapsed together) between the two experimental groups. To ensure that our results on the learning measures were not due to the differences in the average RTs, we repeated the analysis with standardized scores. To this aim, we divided the learning scores (median RTs for low-probability triplets *minus* median RTs for high-probability triplets) by the average of RTs for the high and low-probability triplets of the given epoch for each participant and each epoch.

To see whether participants developed conscious knowledge about the learned statistical regularities, we compared the percentage of the generated high-probability triplets in the Inclusion-Exclusion Task to chance level (25%) separately for the two groups with one-sample t-tests. To reveal if the level of explicitness differs between groups and conditions, we compared the percentage of high-probability triplets with a 2 (Condition: Inclusion vs. Exclusion) × 2 (Group: Accuracy Group vs. Speed Group) mixed-design ANOVA.

Additionally, we correlated the average RTs and accuracy scores with the rates of the different items of the questionnaire to check whether the subjective preferences of the participant are related to the ability to follow the instructions.

## Results

### Did the two groups perform equally before learning?

To ensure that there are no pre-existing differences between groups in terms of speed or accuracy, we compared the median RTs (only for correct responses) and the accuracy of the two groups in the practice session. We did not find differences between groups either in RTs, *t*(59) = -0.48, *p* = .64, or in accuracies, *t*(59) = -1.08, *p* = .28. Therefore, we assumed that there are no pre-existing differences between groups regarding their speed or accuracy.

### Did the instruction affect general RTs and accuracies?

We ran a mixed-design ANOVA with Epoch (1-4) as within-subject factor and Group (Accuracy Group vs. Speed Group) on the average RT and accuracy scores (irrespectively of the stimulus type) to see how the instruction affected the general speed and accuracy of the participants. On RTs measures, a main effect of Epoch was found, F(1.97, 116.33) = 7.46, p = .001, *η*_p_^2^ = .11, indicating a decrease of RTs throughout the task. The main effect of Group was also significant, *F*(1, 59) = 51.84, *p* < .001, *η*_p_^2^ = .47, indicating faster overall RTs of the Speed Group. The Epoch × Group interaction was approaching significance, *F*(3, 177) = 2.30, *p* = .08, *η*_p_^2^ = .04, revealing that on trend level, the rate of RT-decrease was higher in the Speed than in the Accuracy Group (Figure 2).

**Figure 2.**
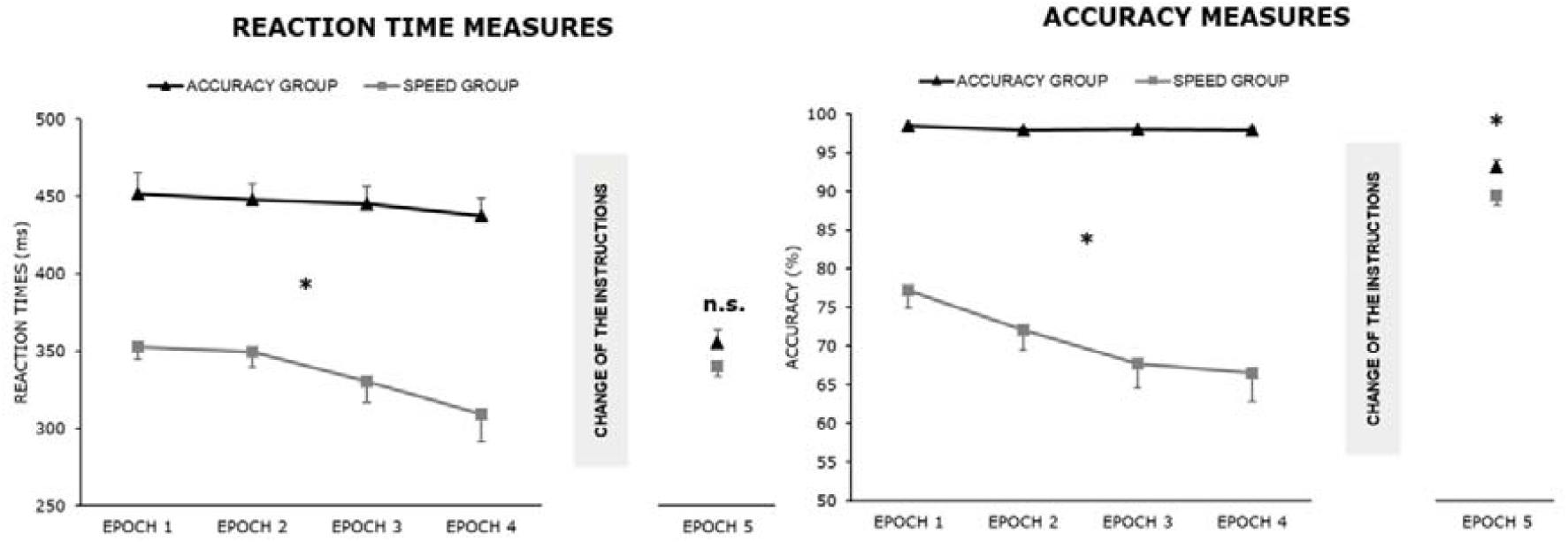
The effects of instruction on average reaction times and accuracies. The horizontal axis indicates the five epochs of the task and the vertical axis the RTs in milliseconds/accuracies in percentage. The error bars represent the standard error. Average RTs were significantly shorter and accuracies lower for the Speed Group from the first epoch, indicating that the participants followed the instructions. After the change of the instructions (Epoch 5) - although the average scores of the two groups approached each other – the difference persisted for accuracies; however, the difference disappeared for the RTs. *.*p* < .05

On accuracy measures, we found a main effect of Epoch, *F*(1.81, 107.10 = 8.19, *p* < 0.001, *η*_p_^2^ = 0.125, revealing a decrease of accuracy as the task progressed. The main effect of Group was also significant, *F*(1, 59) = 117.60, *p* < 0.001, *η*_p_^2^ = 0.67, signaling an increased average accuracy in the Accuracy Group. The Epoch × Group interaction was significant, *F*(3, 177) = 6.91, *p* < 0.001, *η*_p_^2^ = 0.11, indicating that accuracy decreased over the epochs in the Speed Group, while it remained similarly high over all epochs in the Accuracy Group.

### Did the learning process differ between groups as a result of the different instructions?

Next, we investigated whether the sequence learning process differed between groups during the Different Instruction Phase. RTs were analyzed with a mixed-design ANOVA with Triplet (high- vs. low-probability triplets) and Epoch (Epoch 1 to 4) as within-subject factors, and with Group (Accuracy Group vs. Speed Group) as a between-subject factor. The main effect of Triplet was significant, *F*(1, 59) = 49.41, *p* < .001, *η*_p_^2^ = .46: faster RTs were found for high-probability triplets compared to low-probability triplets, revealing implicit statistical learning. Importantly, the Triplet × Group interaction was non-significant, *F*(1, 59) = 0.48, *p* = .49, *η*_p_^2^ = .01, BF_10_ = 0.32): the degree of learning did not differ between the two groups over the course of the learning (Figure 3). The Triplet × Epoch interaction was significant, *F*(3, 177) = 5.66, *p* < .001, *η*_p_^2^ = .09: In the first epoch, no difference was detected between high- and low-probability triplets, while learning (faster RTs for high- than for low-probability triplets) emerged from the second epoch. No difference was found in the time course of statistical learning across groups, as revealed by a non-significant Triplet × Epoch × Group interaction, *F*(3, 177) = 0.90, *p* = .44, *η*_p_^2^ = .02.

**Figure 3.**
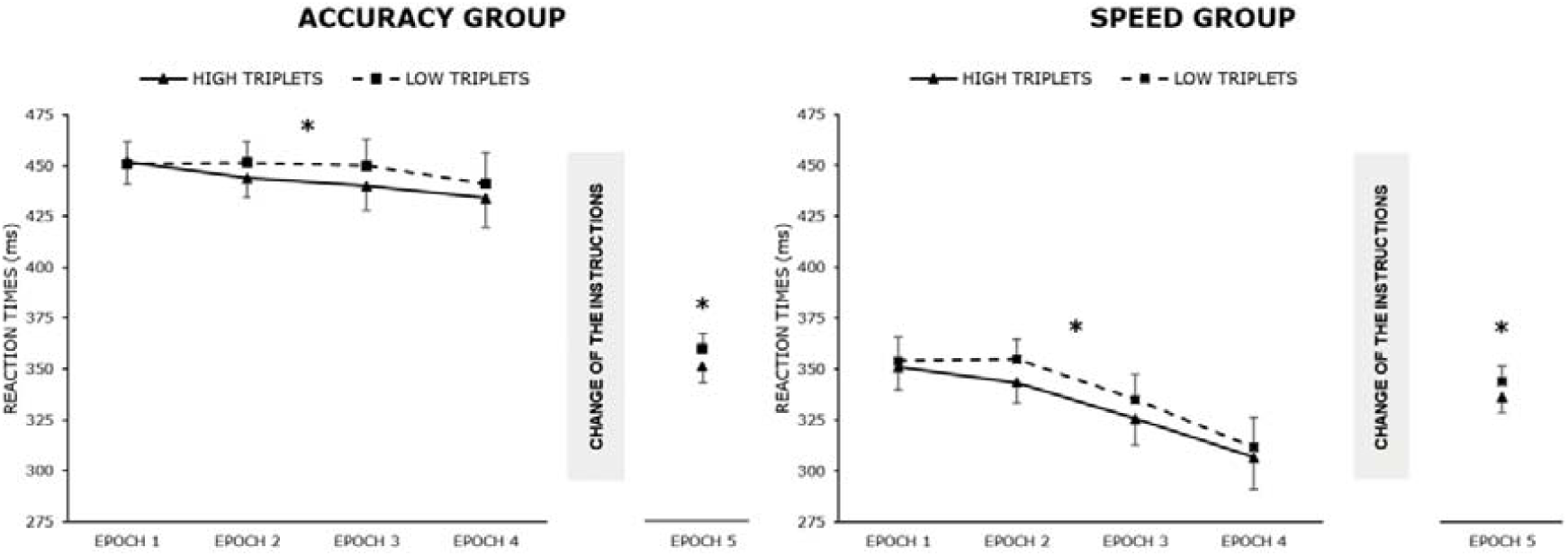
The learning in the two groups. The horizontal axis shows the five epochs of the task and the vertical axis RTs. The dashed line represents the RTs for the high-probability triplets, while the solid line indicates the RTs for the low-probability triplets. The error bars represent the standard error. Please note the learning of statistical regularities is measured by the gap of the two lines. The RTs for high-probability triplets were smaller for both groups and phases. The difference between the two trial types remained after the change of the instructions. A similar level of learning was measured in both groups and phases. *\*.p < .05*

We compared the standardized learning scores between the two groups in each epoch. To this end, a mixed-design ANOVA was performed on the standardized RT learning scores with Epoch (Epoch 1 to 4) as a within-subject factor and Group (Accuracy Group vs. Speed Group) as a between-subject factor. It revealed a main effect of Epoch, *F*(2.09, 123.10) = 2.99, *p* = .048, *η*_p_^2^ = .05, suggesting that, in accordance with the non-standardized data, learning scores did change throughout the task, as the learning scores became larger. The main effect of Group was non-significant, *F*(1, 59) < 0.001, *p* = .53, *η*_p_^2^ = .07, BF_10_ = 0.30, indicating that – in accordance with the non-standardized data – that the two groups exhibited similar learning scores in the task. The interaction of Epoch and Group did not reach significance, *F*(3, 177) = 1.39, *p* = .25, *η*_p_^2^ = .02, indicating the lack of significant group differences in the dynamics of learning over the epochs.

### Did the acquired knowledge on the ASRT task differ between groups when testing with the same instructions?

First, we calculated the median RTs separately for the high- and low-probability triplets at the Similar Instruction Phase. We analyzed RTs of Epoch 5 with a mixed-design ANOVA with Triplet (high-probability triplets vs. low-probability triplets) as within-subject factor and with Group (Accuracy Group vs. Speed Group) as a between-subject factor. A significant main effect of Triplet was revealed, *F*(1, 59) = 50.50, *p* < .001, *η*_p_^2^ = .46, indicating the emergence of acquired statistical knowledge (as RTs on high-probability triplets were smaller than RTs on low-probability triplets). The main effect of Group did not reach significance, *F*(1, 59) = 2.21, *p* = .16, *η*_p_^2^ = .03, indicating that after the change of the instructions, the overall RT difference between groups disappeared (Figure 2). The interaction of Triplet and Group did not reach significance, *F*(1, 59) = 0.27, *p* = .60, *η*_p_^2^ = .01, BF_10_ = 0.29, indicating that irrespectively of the instruction during the learning, the two groups showed the same level of statistical knowledge when no specific instructions were given on the importance of accuracy or speed (Figure 4). Although no difference was found between groups in terms of the average RTs in the Similar Instruction Phase, for the sake of completeness, we repeated the analysis with standardized learning scores. Again, no difference was found between groups in the last epoch, *t*(59) = 0.58, *p* = .57, BF_10_ = 0.30.

**Figure 4.**
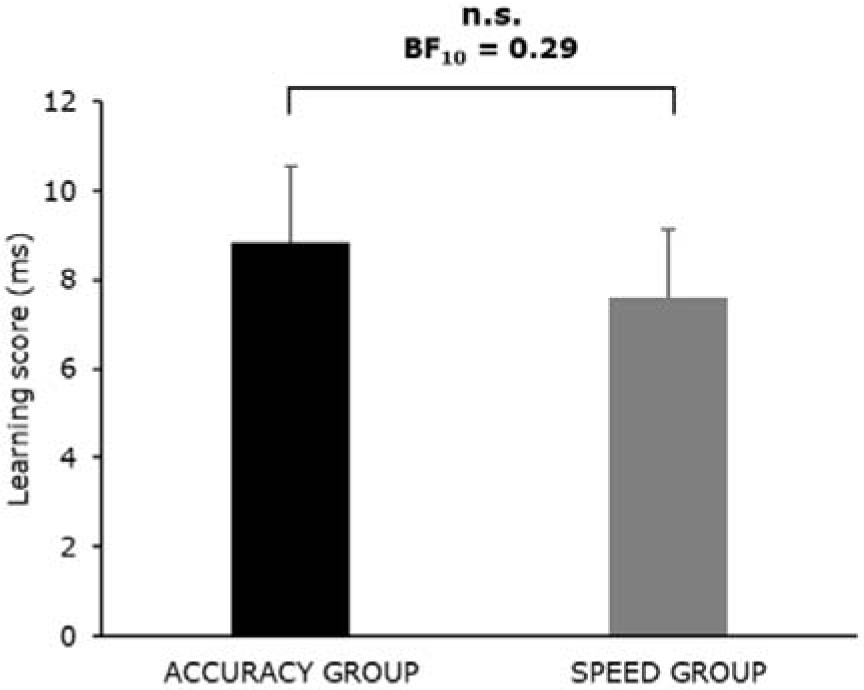
The comparison of learning scores in the Similar Instruction Phase. The vertical axis indicates the learning scores (RTs for low-probability triplets minus RTs for high-probability triplets), and the horizontal axis represents the two groups. The error bars represent the standard error. No significant difference was found between groups, and the lack of difference was confirmed by Bayesian analysis.

### Did the instructions affect the learning if the sequence learning index is based on accuracies?ű

We ran the same Epoch × Triplet × Group mixed-design ANOVA for the Different Instruction Phase and the same Triplet × Group mixed-design ANOVA for the Similar Instruction Phase with accuracy scores. The average accuracies were higher for the Accuracy Group than the Speed Group in the Different Instruction Phase (main effect of Group: *F*(1, 59) = 117.40, *p* < .001, *η*_p_^2^ = .67), and this difference persisted after the change of the instructions (main effect of Group: *F*(1, 59) = 5.08, *p* = .03, *η*_p_^2^ = .08) (Figure 2). In the Different Instruction Phase, we found evidence for statistical knowledge in the Speed Group (the high- and low-probability triplets differed from each other), while the Accuracy Group did not show evidence for learning (Triplet × Group: *F*(1, 59) = 45.25, *p* < .001, *η*_p_^2^ = .43). However, we did not find difference between groups in terms of the acquired knowledge when no speed/accuracy instructions were given, *F*(1, 59) = 0.85, *p* = .36, *η*_p_^2^ = .01, BF_10_ = 0.37. The same results were found with standardized scores. The details of the analysis and the results can be found in the Supplementary Material.

### Did the participants develop conscious knowledge about the statistical regularities, and was it different between groups?

The Inclusion/Exclusion task was administered to reveal whether the acquired statistical knowledge remained implicit or became explicitly accessible for the participant. Separately for the two groups, we compared the percentage of the generated high-probability triplets to the chance level (25%). In the Accuracy Group, two participants were excluded from this analysis as they did not follow the instructions. Participants in the Accuracy Group generated 32.33% high-probability triplets in the Inclusion condition, which is significantly higher than chance level, *t*(28) = 4.82, *p* < .001. Participants of the Accuracy Group generated high-probability triplets significantly above chance (29.81%) in the Exclusion condition as well *t*(28) = 4.04, *p* = .001, indicating that they could not consciously control the emergence of this knowledge. In the Speed Group, two participants were excluded as they did not follow the instructions. Participants of the Speed Group generated 29.34% high-probability triplets in the Inclusion condition, which is significantly higher than chance level, *t*(27) = 3.58, *p* = .001. They also generated more high-probability triplets than expected by chance in the Exclusion condition, 29.25%; *t*(27) = 2.07, *p* = .048.

Furthermore, we compared the differences between groups and between tasks with a 2 (Condition: Inclusion vs. Exclusion) × 2 (Group: Accuracy Group vs. Speed Group) ANOVA. The main effect of Condition was not significant, *F*(1, 55) = 1.66, *p* = .20, *η*_p_^2^ = .03, thus, the triplet knowledge of the participants remained implicit. The Group main effect did not reach significance, *F*(1, 55) = 0.53, *p* = .47, *η*_*p*_^*2*^ = .01, indicating that the two groups performed equally on the two tasks. The interaction of the Condition and the Group factors was not significant, *F*(1, 55) = 0.26, *p* = .61, *η*_p_^2^ = .01, meaning that the lack of difference between groups was not influenced by the type of the task.

### Did the preferences of the participants affect their performance on the task?

We used a questionnaire to check whether the subjective preferences on being fast or accurate in the real-life were related to the ability to follow instructions (see Methods for the questions). We correlated the questionnaire scores with the average RTs and accuracy of the participants separately for the two groups. We did not find any significant correlations between the average scores and subjective ratings about the preferences either in the Accuracy Group or in the Speed Group (all *p* > .01). It indicates that the preference of accuracy or speed, and whether the participants are rather fast or accurate in real life did not play a role in the ability to follow the instructions.

## Discussion

Here, we aimed to unveil whether speed/accuracy instructions can influence an essential compound of skill learning, namely the acquisition of probabilistic statistical regularities learning. To this end, we instructed two groups of participants to be either fast or accurate during the training in our implicit probabilistic sequence learning task. In the testing phase, we assessed the accumulated knowledge of probabilistic regularities, and this time, all of the participants were instructed to be equally fast and accurate. As predicted, the instructions greatly affected the general speed and accuracy of the participants: the speed instructions resulted in faster reaction times and a higher number of errors, while the accuracy instructions caused slower overall reaction times and an almost errorless performance. Despite these differences during training, the sequence learning index based on RTs was similar in both groups. Thus, the instructions did not affect the acquisition of implicit probabilistic regularities during the training. Moreover, no difference between the groups was found in the testing phase. This lack of difference suggests that instructions did not affect either the performance during training or the acquired statistical knowledge. Similar results were obtained when we controlled for the differences in average speed between groups. Moreover, Bayesian statistical methods also supported the lack of difference between groups in terms of the acquired knowledge.

Our main result is that we detected a similar level of acquired knowledge irrespective of the strategy used during the training. This finding has several implications. From a narrower, learning perspective, it suggests that our ability to extract the relevant pieces of statistical information from the environment is so robust that instructions cannot influence it. It is in accordance with the findings that statistical knowledge persists and remains resistant to interference even after one-year (Kóbor et al., 2017), is intact in dual-task conditions (Vékony et al., 2019) or in certain disorders characterized by cognitive dysfunctions, such as obstructive sleep apnea (Csabi, Varszegi-Schulz, Janacsek, Malecek, & Nemeth, 2014; Nemeth, Csábi, Janacsek, Várszegi, & Mari, 2012), sleep-disordered breathing (Csábi, Benedek, Janacsek, Katona, & Nemeth, 2013; Csábi et al., 2016), autism (Nemeth, Janacsek, Balogh, et al., 2010), borderline personality disorder (Unoka et al., 2017) or alcohol-dependency (Virag et al., 2015). Deterministic learning tasks test patterns that occur with a 100% probability over time, while the alteration of the random and pattern elements in the ASRT task creates a noisy, uncertain environment, which is similar to the natural environments of learning in everyday life (Fiser, Berkes, Orbán, & Lengyel, 2010). Our results showed that using complex probabilistic regularities, a similar level of statistical knowledge emerges throughout learning, even when learning occurs under different circumstances, with different strategies.

Another compelling result of our study is that participants in the accuracy condition did acquire stable statistical knowledge despite the minimization of motor (response) errors during the training. This statistical knowledge was equal to the knowledge acquired with speed instructions, which was characterized by a relatively high amount of errors during the training. This result is especially interesting in the light of the theory claiming that the brain is a Bayesian inference machine (Friston, 2010) because our results contradict the findings that committing errors facilitates learning (Bubic, Von Cramon, & Schubotz, 2010). Our brain learns associations between events through the continuous adjustments of the estimated probability distribution, i.e., the prior. After a prediction error, the prior should be updated in accordance with the new information about the probabilistic structure (Friston, 2010). Based on these theories, we would expect a low number of errors to impair the learning process. This was not the case in our study, which raises the possibility that the motor aspect of prediction errors is not crucial in all circumstances for updating the priors during probabilistic sequence learning. However, it is also possible that a similar amount of errors might be detected with other methods, for example, by investigating eye-movements (Le Pelley, Beesley, & Griffiths, 2011; Wills, Lavric, Croft, & Hodgson, 2007). The exploration of the role of errors in implicit sequence learning deserves future investigation, using eye-tracking during sequence learning and electrophysiological methods to measure error-related brain activity.

At the initial training phase, a similar level of statistical learning was found under speed and accuracy instructions. The fact that no group differences were found is in contrast with the results of Hoyndorf and Haider (2009), as they detected impaired implicit performance with accuracy strategy. In their study, participants performed a regular and a random task set during a Number Reduction Task. They found that only the participants focusing on speed had increased speed for the regular task set. The authors claimed that the increased monitoring due to the accuracy instruction might have impeded the performance, similarly as in skill acquisition studies (Beilock et al., 2008, 2004). However, in the same study, Hoyndorf and Haider (2009) found a preference for the regular task set also in the accuracy group, which they interpreted as that the focus on accuracy affects only the expression of implicitly acquired knowledge rather than learning processes per se. This is in accordance with our results, as we found a similar level of sequence knowledge when we equally emphasized the importance of speed and accuracy after the initial learning. The difference regarding the training phase might be explained by the fact that we studied more complex, probabilistic sequence representations, which might be more resistant to speed and accuracy strategies than deterministic patterns. Similarly, Barnhoorn et al (2019), who have also found speed instruction to benefit the development of sequence representations, used simple repeating sequences. Moreover, this study investigated explicit sequence learning processes, while our participants were unaware of their accumulated sequence knowledge. A possible explanation for the difference between the effect of implicit and explicit learning conditions could be that the increased speed covers up the explicitness of the task. As a consequence, the task becomes more implicit, the top-down control reduces, and the learning becomes better. In our study, the learning was entirely implicit, therefore, the speeding up could not improve the level of implicitness. Thus, the learning was similar under speed and accuracy instructions. Future investigation is needed to reveal to what extent is the implicit or the probabilistic nature of the task related to lack of speed benefit during training.

From a broader, cognitive neuroscience perspective, it is essential to highlight the relationship between learning and performance in our study. Most studies in the field of cognitive neuroscience measure learning at one time point, and draw conclusions about brain-behavior relationships based on either the *long-term learning* (the relatively permanent changes in knowledge, i.e., competence) or the *momentary performance* (the temporary fluctuation in behavior) (e.g., Heideman, van Ede, & Nobre, 2018; Rose, Haider, Salari, & Buchel, 2011; Thomas et al., 2004; Turk-Browne, Scholl, Johnson, & Chun, 2010). However, it was shown that these two factors could be separated from each other. For example, learning and performance can differ due to fatigue, different types of practice, latent learning, or overlearning of the practiced skill (Soderstrom & Bjork, 2015). Our study also revealed that the learning of a certain skill could differ from the momentary performance due to different instructions, at least when the accuracy is used as an indicator (see Supplementary Material). This result draws attention to the problem of using solely one session to evaluate learning. For example, if fatigue or boredom of the participants are different when they concentrate on being fast or accurate, then it can influence the conclusions we draw from our results. However, when the learning index (difference score) is based on RTs, this contingency appears smaller, at least when investigating implicit probabilistic sequence learning. Future studies should reveal to what extent this phenomenon is generalizable to other types of learning, such as to more explicit or to non-statistical learning tasks. Non-learning tasks should also be tested, as general speed-up and changes in accuracy can be seen over the course of several cognitive tasks requiring fast decision-making. Based on our results, we recommend taking into consideration the possible differences between the measured competence and performance when designing learning studies.

We manipulated the general speed and accuracy of the participants by giving explicit instructions to focus either on speed or accuracy, as previous non-learning cognitive tasks also did (e.g., Aasen & Brunner, 2016; Christensen et al., 2001; Osman et al., 2000; Ullsperger, Bylsma, & Botvinick, 2004). However, it might be questionable if our results truly reflect the effect of instructions on learning. One can argue that the instructions given in our study were not strong enough to manipulate the learning strategy and therefore, the learning processes because previous studies on the topics used more pronounced instructions and signals to modify the strategy of the participants (Barnhoorn et al., 2019; Hoyndorf & Haider, 2009). This seems unlikely as the general speed and accuracy were affected by the instructions. Group differences also emerged in *general skill learning* as (1) participants who focused on their speed showed increasingly faster responses, and (2) participants who focused on their accuracy sustained a high level of accuracy during the learning phase compared to the other group. In contrast to these findings, the acquisition of statistical regularities was not affected by the instructions. To sum up, we found evidence that speed and accuracy affect general skill learning and sequence-specific learning (statistical learning) differently.

It can also be claimed that verbal instructions given at the beginning of the task might not be sufficient to regulate subjects’ average speed and accuracy, because as time goes on, participants tend to wane in favor of their response tendencies (Heitz, 2014). In other words, they will behave according to their preferences for being accurate or fast on a task. In our case, this change in behavior is unlikely. First, we found no differences in the average RTs and accuracy scores between groups when the participants practiced the task on random sequences (before we gave distinct instructions to the groups), and second, participants did not become less accurate or slower throughout the task. Therefore, the effects observed should be the results of the instructions. Additionally, we measured the participants’ individual preferences on response tendencies with a questionnaire (whether they prefer to be accurate or fast). No correlations were observed between these individual preferences and the average speed and accuracy during the task in either group. These aspects indicate that our results indeed reflect the effect of instructions, and participants did not follow their individually preferred response tendencies during the task.

## Conclusions

Our study investigated the effects of speed and accuracy instructions on an essential compound of skill learning, namely the acquisition of probabilistic regularities. Our main finding is that our ability to pick-up statistical regularities in a noisy, uncertain environment is so robust that instructions do not influence it. It indicates that implicit probabilistic sequence learning is independent of the manipulation of speed/accuracy trade-off. Another finding of our study is that the learning is intact with almost 100% accuracy level. It suggests that statistical learning is at least partly independent of accuracy level, and statistical knowledge about the environmental regularities can be acquired even if no response (motor) errors occur. Our results also raise the possibility that competence and performance can differ in some instances. Accuracy instructions can mask the accumulating knowledge during learning when measured by accuracy, although statistical knowledge does emerge in these cases as well. Future studies investigating whether this robustness is related to the implicit feature of the task or whether different types of learning are affected equally, seem warranted.

## Acknowledgment

This research was supported by the National Brain Research Program (project 2017-1.2.1-NKP-2017-00002); Hungarian Scientific Research Fund (NKFIH-OTKA K 128016, PI: D. N., NKFIH-OTKA PD 124148, PI: K.J.); Janos Bolyai Research Fellowship of the Hungarian Academy of Sciences (to K. J. and to A. M.); EFOP-3.6.1-16-2016-00008 (to A. M.); IDEXLYON Fellowship of the University of Lyon as part of the Programme Investissements d’Avenir (ANR-16-IDEX-0005) (to D.N). The authors are grateful to Lúcia Nemes, Soma Béres, and Réka Sefcsik for their help in data acquisition.

## References

Aasen, I. E., & Brunner, J. F. (2016). Modulation of ERP components by task instructions in a cued go/no-go task. Psychophysiology, 53(2), 171–185. https://doi.org/10.1111/psyp.12563

Barnhoorn, J. S., Panzer, S., Godde, B., & Verwey, W. B. (2019). Training motor sequences: Effects of speed and accuracy instructions. Journal of Motor Behavior, 51(5), 540–550. https://doi.org/10.1080/00222895.2018.1528202

Beilock, S. L., Bertenthal, B. I., Hoerger, M., & Carr, T. H. (2008). When does haste make waste? Speed-Accuracy tradeoff, skill level, and the tools of the trade. Journal of Experimental Psychology: Applied, 14(4), 340–352. https://doi.org/10.1037/a0012859

Beilock, S. L., Bertenthal, B. I., McCoy, A. M., & Carr, T. H. (2004). Haste does not always make waste: Expertise, direction of attention, and speed versus accuracy in performing sensorimotor skills. Psychonomic Bulletin and Review, 11(2), 373–379. https://doi.org/10.3758/BF03196585

Bubic, A., Von Cramon, D. Y., & Schubotz, R. I. (2010). Prediction, cognition and the brain. Frontiers in Human Neuroscience, 4(25), 1–15. https://doi.org/10.3389/fnhum.2010.00025

Christensen, C. A., Ivkovich, D., & Drake, K. J. (2001). Late positive ERP peaks observed in stimulus-response compatibility tasks tested under speed-accuracy instructions. Psychophysiology, 38(3), 404–416. https://doi.org/10.1017/S0048577201991176

Christiansen, M. H., & Chater, N. (2015). The language faculty that wasn’t: a usage-based account of natural language recursion. Frontiers in Psychology, 6, 1182. https://doi.org/10.3389/fpsyg.2015.01182

Conway, C. M. (2020). How does the brain learn environmental structure? Ten core principles for understanding the neurocognitive mechanisms of statistical learning. Neuroscience and Biobehavioral Reviews, 112, 279–299. https://doi.org/10.1016/j.neubiorev.2020.01.032

Csábi, E., Benedek, P., Janacsek, K., Katona, G., & Nemeth, D. (2013). Sleep disorder in childhood impairs declarative but not nondeclarative forms of learning. Journal of Clinical and Experimental Neuropsychology, 35(7), 677–685. https://doi.org/10.1080/13803395.2013.815693

Csábi, E., Benedek, P., Janacsek, K., Zavecz, Z., Katona, G., & Nemeth, D. (2016). Declarative and non-declarative memory consolidation in children with sleep disorder. Frontiers in Human Neuroscience, 9(709). https://doi.org/10.3389/fnhum.2015.00709

Csabi, E., Varszegi-Schulz, M., Janacsek, K., Malecek, N., & Nemeth, D. (2014). The consolidation of implicit sequence memory in obstructive sleep apnea. PLoS ONE, 9(10), 1–6. https://doi.org/10.1371/journal.pone.0109010

Destrebecqz, A., & Cleeremans, A. (2001). Can sequence learning be implicit? New evidence with the process dissociation procedure. Psychonomic Bulletin and Review, 8(2), 343–350. https://doi.org/10.3758/BF03196171

Destrebecqz, A., Peigneux, P., Laureys, S., Degueldre, C., Del Fiore, G., Aerts, J., … Maquet, P. (2005). Neural correlates of implicit and explicit sequence learning: Interacting networks revealed. Learning & Memory, 12(5), 480–490. https://doi.org/10.1101/lm.95605.6

Fiser, J., Berkes, P., Orbán, G., & Lengyel, M. (2010). Statistically optimal perception and learning: from behavior to neural representations. Trends in Cognitive Sciences, 14(3), 119–130. https://doi.org/10.1016/j.tics.2010.01.003

Friston, K. (2010). The free-energy principle: A unified brain theory? Nature Reviews Neuroscience, 11(2), 127–138. https://doi.org/10.1038/nrn2787

Fu, Q., Dienes, Z., & Fu, X. (2010). Can unconscious knowledge allow control in sequence learning? Consciousness and Cognition, 19(1), 462–474. https://doi.org/10.1016/j.concog.2009.10.001

Heideman, S. G., van Ede, F., & Nobre, A. C. (2018). Temporal alignment of anticipatory motor cortical beta lateralisation in hidden visual-motor sequences. European Journal of Neuroscience, 48(8), 2684–2695. https://doi.org/10.1111/ejn.13700

Heitz, R. P. (2014). The speed-accuracy tradeoff: History, physiology, methodology, and behavior. Frontiers in Neuroscience, 8(150), 1–19. https://doi.org/10.3389/fnins.2014.00150

Horvath, K., Torok, C., Pesthy, O., Nemeth, D., & Janacsek, K. (2018). Explicit instruction differentially affects subcomponents of procedural learning and consolidation. BioRxiv, 433243. https://doi.org/10.1101/433243

Howard, J. H., & Howard, D. V. (1997). Age differences in implicit learning of higher order dependencies in serial patterns. Psychology and Aging, 12(4), 634–656. https://doi.org/10.1037/0882-7974.12.4.634

Howard, D. V., Howard, J. H., Japikse, K., DiYanni, C., Thompson, A., & Somberg, R. (2004). Implicit sequence learning: Effects of level of structure, adult age, and extended practice. Psychology and Aging, 19(1), 79–92. https://doi.org/10.1037/0882-7974.19.1.79

Hoyndorf, A., & Haider, H. (2009). The “Not Letting Go” phenomenon: Accuracy instructions can impair behavioral and metacognitive effects of implicit learning processes. Psychological Research, 73(5), 695–706. https://doi.org/10.1007/s00426-008-0180-4

Jacoby, L. L. (1991). A process dissociation framework: Separating automatic from intentional uses of memory. Journal of Memory and Language, 30(5), 513–541.

Janacsek, K., Borbély-Ipkovich, E., Nemeth, D., & Gonda, X. (2018). How can the depressed mind extract and remember predictive relationships of the environment? Evidence from implicit probabilistic sequence learning. Progress in Neuro-Psychopharmacology and Biological Psychiatry, 81, 17–24. https://doi.org/10.1016/j.pnpbp.2017.09.021

Janacsek, K., Fiser, J., & Nemeth, D. (2012). The best time to acquire new skills: Age-related differences in implicit sequence learning across the human lifespan. Developmental Science, 15(4), 496–505. https://doi.org/10.1111/j.1467-7687.2012.01150.x

JASP Team. (2019). JASP (Version 0.10).

Jiménez, L., Vaquero, J. M. M., & Lupiáñez, J. (2006). Qualitative differences between implicit and explicit sequence learning. Journal of Experimental Psychology: Learning Memory and Cognition, 32(3), 475–490. https://doi.org/10.1037/0278-7393.32.3.475

Kiss, M., Nemeth, D., & Janacsek, K. (2019). Stimulus presentation rates affect performance but not the acquired knowledge – Evidence from procedural learning. BioRxiv, 650598.

Kóbor, A., Janacsek, K., Takács, A., Nemeth, D., Kobor, A., Janacsek, K., … Nemeth, D. (2017). Statistical learning leads to persistent memory: Evidence for one-year consolidation. Scientific Reports, 7(1), 1–10. https://doi.org/10.1038/s41598-017-00807-

Le Pelley, M. E., Beesley, T., & Griffiths, O. (2011). Overt attention and predictiveness in human contingency learning. Journal of Experimental Psychology: Animal Behavior Processes, 37(2), 220–229. https://doi.org/10.1037/a0021384

Nemeth, D., Csábi, E., Janacsek, K., Várszegi, M., & Mari, Z. (2012). Intact implicit probabilistic sequence learning in obstructive sleep apnea. Journal of Sleep Research, 21(4), 396–401. https://doi.org/10.1111/j.1365-2869.2011.00983.x

Nemeth, D., Janacsek, K., Balogh, V., Londe, Z., Mingesz, R., Fazekas, M., … Vetro, A. (2010). Learning in autism: Implicitly superb. PLoS ONE, 5(7), 1–7. https://doi.org/10.1371/journal.pone.0011731

Nemeth, D., Janacsek, K., Király, K., Londe, Z., Németh, K., Fazekas, K., … Csányi, A. (2013). Probabilistic sequence learning in mild cognitive impairment. Frontiers in Human Neuroscience, 7, 318. https://doi.org/10.3389/fnhum.2013.00318

Nemeth, D., Janacsek, K., Londe, Z., Ullman, M. T., Howard, D. V., & Howard, J. H. (2010). Sleep has no critical role in implicit motor sequence learning in young and old adults. Experimental Brain Research, 201(2), 351–358. https://doi.org/10.1007/s00221-009-2024-x

Oldfield, R. C. (1971). The assessment and analysis of handedness: The Edinburgh inventory. Neuropsychologia, 9(1), 97–113. https://doi.org/10.1016/0028-3932(71)90067-4

Osman, A., Lou, L., Muller-Gethmann, H., Rinkenauer, G., Mattes, S., & Ulrich, R. (2000). Mechanisms of speed-accuracy tradeoff: Evidence from covert motor processes. Biological Psychology, 51(2–3), 173–199. https://doi.org/10.1016/S0301-0511(99)00045-9

Remillard, G. (2008). Implicit learning of second-, third-, and fourth-order adjacent and nonadjacent sequential dependencies. Quarterly Journal of Experimental Psychology, 61(3), 400–424. https://doi.org/10.1080/17470210701210999

Rose, M., Haider, H., Salari, N., & Buchel, C. (2011). Functional dissociation of hippocampal mechanism during implicit learning based on the domain of associations. Journal of Neuroscience, 31(39), 13739–13745. https://doi.org/10.1523/jneurosci.3020-11.2011

Soderstrom, N. C., & Bjork, R. A. (2015). Learning versus performance: An integrative review. Perspectives on Psychological Science, 10(2), 176–199. https://doi.org/10.1177/1745691615569000

Song, S., Howard, J. H., & Howard, D. V. (2007). Sleep does not benefit probabilistic motor sequence learning. Journal of Neuroscience, 27(46), 12475–12483. https://doi.org/10.1523/jneurosci.2062-07.2007

Takács, Á., Kóbor, A., Chezan, J., Éltető, N., Tárnok, Z., Nemeth, D., … Janacsek, K. (2018). Is procedural memory enhanced in Tourette syndrome? Evidence from a sequence learning task. Cortex, 100, 84–94. https://doi.org/10.1016/j.cortex.2017.08.037

Thomas, K. M., Hunt, R. H., Vizueta, N., Sommer, T., Durston, S., Yang, Y., & Worden, M. S. (2004). Evidence of developmental differences in implicit sequence learning: An fMRI study of children and adults. Journal of Cognitive Neuroscience, 16(8), 1339–1351. https://doi.org/10.1162/0898929042304688

Turk-Browne, N. B., Scholl, B. J., Johnson, M. K., & Chun, M. M. (2010). Implicit perceptual anticipation triggered by statistical learning. Journal of Neuroscience, 30(33), 11177–11187. https://doi.org/10.1523/jneurosci.0858-10.2010

Ullsperger, M., Bylsma, L. M., & Botvinick, M. M. (2004). The conflict-adaptation effectC: it ‘s not just priming. Cognitive, Affective, & Behavioral Neuroscience, 5(4), 467–472.

Unoka, Z., Vizin, G., Bjelik, A., Radics, D., Nemeth, D., & Janacsek, K. (2017). Intact implicit statistical learning in borderline personality disorder. Psychiatry Research, 255, 373–381. https://doi.org/10.1016/j.psychres.2017.06.072

Vékony, T., Török, L., Pedraza, F., Schipper, K., Plèche, C., Tóth, L., … Nemeth, D. (2019). Retrieval of a well-established skill is resistant to distraction: evidence from an implicit probabilistic sequence learning task. BioRxiv, 849729. https://doi.org/10.1101/849729

Virag, M., Janacsek, K., Horvath, A., Bujdoso, Z., Fabo, D., & Nemeth, D. (2015). Competition between frontal lobe functions and implicit sequence learning: evidence from the long-term effects of alcohol. Experimental Brain Research, 233(7), 2081–2089. https://doi.org/10.1007/s00221-015-4279-8

Wills, A. J., Lavric, A., Croft, G. S., & Hodgson, T. L. (2007). Predictive learning, prediction errors, and attention: Evidence from event-related potentials and eye tracking. Journal of Cognitive Neuroscience, 19(5), 843–854. https://doi.org/10.1162/jocn.2007.19.5.843

