## Supplementary Material for "Skill learning can be independent of speed and accuracy instructions"

**Author Note**

Correspondence concerning this article should be addressed to Dezso Nemeth, Lyon Neuroscience Research Center (CRNL), INSERM, CNRS, Université de Lyon, Centre Hospitalier Le Vinatier - Bâtiment 462 - Neurocampus 95 boulevard Pinel 69675 Bron, France. Phone: +33 4 81 10 65 46

### Analysis of statistical learning on accuracy measures

Besides the analysis of the reaction times, we performed the complete analysis based on accuracies too. To investigate whether learning process differed between groups during the Different Instruction Phase, accuracies were analyzed with mixed-design ANOVAs with Triplet (high vs. low-probability triplets) and Epoch (Epoch 1 to 4) as within-subject factors, and Group (Accuracy Group vs. Speed Group) as between-subject factor. To check if there was a difference between groups in the Similar Instruction Phase, we analyzed accuracies of Epoch 5 with mixed-design ANOVAs with Triplet (high- vs. low-probability triplets) within-subject factor and the Group (Accuracy Group vs. Speed Group) between-subject factor.

To correct for the average accuracy difference between groups caused by the instructions, we divided the learning scores (mean accuracies for high-probability triplets *minus* low-probability triplets) by the accuracy of the given epoch for each participant and each epoch. For the Different Instruction Phase, mixed-design ANOVAs were performed on the standardized learning scores with the Epoch (Epoch 1 to 4) within-subject factor and the Group (Accuracy Group vs. Speed Group) between-subject factor. For the Similar Instruction Phase, independent samples t-tests were performed on the standardized learning scores between the two experimental groups.

### Results

#### Did the training performance measured by accuracy differ between groups as a result of the different instructions?

We compared mean accuracy scores between the two groups in terms of epochs and triplet types. The main effect of Triplet was significant, *F*(1, 59) = 93.88, *p* < .001*,* *η*_p_^2^ = .65: participants were more accurate responding to high-frequency triplets compared to the low-frequency probability ones, revealing implicit statistical learning. Contrary to the RT results, the Triplet × Group interaction was significant, *F*(1, 59) = 45.25, *p* < .001, *η*_p_^2^ = .43. While the Speed Group showed more accurate responses to high-probability triplets compared to the low-probability ones, the Accuracy Group exhibited similarly accurate responses to high- and low-probability triplets. The Triplet × Epoch interaction proved to be significant, *F*(3, 177) = 3.69, *p* = .01, *η*_p_^2^ = .06: participants' accuracy for low-probability triplets decreased over the course of training, while it remained relatively constant for high-probability triplets, which reflects increasing implicit statistical learning. The Epoch × Triplet × Group interaction was also significant, *F*(3, 177) = 2.987, *p* = .03, *η*_p_^2^ = .05, suggesting different dynamics of implicit statistical learning for the two groups, with increasingly greater learning in the Speed Group compared to the Accuracy Group.

We also compared the standardized accuracy learning scores between the two groups in each epoch. The Epoch × Group ANOVA on the corrected learning scores revealed a significant main effect of Epoch, *F*(3, 177) = 5.21, *p* = .002, *η*_p_^2^ = .08, such that learning scores increased over the course of the task. Importantly, the main effect of Group was significant again, *F*(1, 59) = 46.17, *p* < .001, *η*_p_^2^ = .44: the Speed Group showed accuracy-related learning, while the Accuracy Group did not. The Epoch × Group interaction was also significant, *F*(3, 177) = 4.82, *p* = .003, *η*_p_^2^ = .08, indicating that increasingly greater learning scores only in the Speed Group.

#### Did the acquired knowledge measured by accuracies differ between groups when the importance of accuracy and speed were equally emphasized?

The Triplet × Group ANOVA revealed a significant main effect of Triplet, *F*(1, 59) = 39.96, *p* < .001, *η*_p_^2^ = .40, indicating the acquired knowledge on the task in accuracy as well (more accurate responses for high-probability triplets compared to the low-probability ones). The main effect of Group was significant, *F*(1, 59) = 5.08, *p* = .03, *η*_p_^2^ = .08, indicating that the overall difference in accuracy persisted after the change of the instructions. The Triplet × Group interaction did not reach significance, *F*(1, 59) = 0.85, *p* = .36, *η*_p_^2^ = .01, signaling a similar level of competence on the task after the change of the instruction. We also compared the competence of the two groups with corrected learning scores. In this case, we did not find difference between the two groups either, *t*(59) = -0.89, *p* = .38.
